# Inhibiting the cholesterol storage enzyme ACAT1/SOAT1 in aging Apolipoprotein E4 mice alter their brains inflammatory profiles

**DOI:** 10.1101/2024.10.24.620063

**Authors:** Thao N. Huynh, Emma N. Fikse, Matthew C. Havrda, Catherine C.Y. Chang, Ta Yuan Chang

## Abstract

Aging and Apolipoprotein E4 (APOE4) are the two most significant risk factors for late-onset Alzheimer’s disease (LOAD). Compared to APOE3, APOE4 disrupts cholesterol homeostasis, increases cholesteryl esters (CEs), and exacerbates neuroinflammation in brain cells including microglia. Targeting CEs and neuroinflammation could be a novel strategy to ameliorate APOE4 dependent phenotypes. Toll-like receptor 4 (TLR4) is a key player in inflammation, its regulation is associated with cholesterol content of lipid rafts in cell membranes. We previously demonstrated that in normal microglia expressing APOE3, inhibiting the cholesterol storage enzyme acylCoA:cholesterol acyltransferase 1 (ACAT1/SOAT1) reduces CEs, dampened neuroinflammation via modulating the fate of TLR4. We also showed that treating myelin debris-loaded normal microglia with ACAT inhibitor F12511 reduced cellular CEs and activated ABC transporter 1 (ABCA1) for cholesterol efflux. In this study, we found that treating primary microglia expressing APOE4 with F12511 also reduces CEs, activated ABCA1, and dampened LPS dependent NFkB activation. In vivo, a two-week injections of nanoparticle F12511, which consists of DSPE-PEG_2000_, phosphatidylcholine, and F12511, to aged female APOE4 mice reduced TLR4 protein content and decreased proinflammatory cytokines including IL-1β in APOE4 mice brains. Overall, our work suggests nanoparticle F12511 is a novel agent to ameliorate LOAD.

## 1. Introduction

Alzheimer’s disease (AD) is a progressive form of dementia that significantly reduces patients’ quality of life. Those with AD experience memory loss, cognitive decline, and behavioral changes. Aging is the leading cause for AD and is associated with late disease onset (LOAD) [1]. About 90-95% of AD patients have LOAD [2]. Currently, 1 in 9 people aged 65 years and older have the disease [3]. As people aged, the risk of having AD increases significantly. The *APOE4* gene variants further elevate this risk in carriers between 4 to 15-fold [4,5]. It is not clear how ApoE4 increases the risk for AD. APOE is a protein primarily function in cholesterol and lipid handling in the body and in the brain [5]. APOE has three different isoforms (E2, E3 and E4), in which E2 is associated with lower disease risk, E3 is recognized as associated with the “normal” phenotypes and E4 is associated with increase AD risk [5]. Interestingly, between these three isoforms, there are only different in two amino acids in their sequence that drastically impact their down-stream phenotypes [5]. APOE4 is reported to cause dysfunction across multiple cell types in the central nervous system (CNS), the periphery and the vasculature [6-12].

As originally characterized by Dr. Alois Alzheimer, AD patients typically carry three main pathological hallmarks: the deposition of amyloid beta (Aβ) plaque, tau neurofibrillary tangles (NFT) and the accumulation of lipid granules (Reviewed in [13,14]). These pathological hallmarks in AD had been demonstrated to be associated with neuroinflammation, which correlated with worsen disease progression [15]. Most research in the field has been focusing on tackling Aβ plaque and NFT to ameliorate disease progression and symptoms, with much less focus on lipids [16]. Recently, more advanced techniques allowed researchers to further investigate the role of lipids and cholesterol in disease pathogenesis and progression (Reviewed in [17]).

In recent years, advancement towards developing an Alzheimer’s therapy has made good progress with FDA approvals of a few amyloid targeted immunotherapy. However, recent studies have shown that a few patients carrying *APOE4* gene had died from amyloid-related imaging abnormalities (ARIA). Clinical studies suggest that some AD patients with *APOE4* homozygotes may have an “explosive” response to amyloid immunotherapy [18,19]. Thus, there is an urgent need for an alternative therapy for APOE4 patient population.

In the brain, APOE4 is produced mainly by two cell types: astrocyte and microglia. APOE4 impairs immune response and associates with increasing in lipid droplets in the microglia [10-12] and (Reviewed in [17,20]). Increased lipid droplet accumulation in APOE4 microglia is associated with AD pathology and worsen neuroinflammation [10-12] and (Reviewed in [17,20]).

Lipid droplets contain neutral lipids, predominantly triacylglycerol (TAG) and cholesteryl ester (CE). While most of the recent published studies focused on removing excess TAG to improve APOE4 phenotypes, studies on removing excess CE are still limited (Reviewed in [20]). In cells, CE are synthesized by the cholesterol storage enzyme acyl-CoA: cholesterol acyltransferase, also known as sterol O-acyltransferase 1 (ACAT1/SOAT1). Genetic ablation and pharmaceutical inhibition of ACAT1/SOAT1 have been reported to reduce CE, ameliorated AD associated pathological hallmarks and attenuated neuroinflammation via modulating toll-like receptor 4 (TLR4) [21-28]. In the context of APOE4, pharmaceutical inhibition of ACAT/SOAT using Avasimibe (CI-1011) was reported to be beneficial in 5xFAD/APOE4 mice by reducing intracellular lipid droplets and neuroinflammation, increased memory performance, as well as postsynaptic protein levels [27]. However, whether ACAT1/SOAT1 pharmaceutical inhibition would be beneficial in APOE4 aging model (a model that capture two of the greatest risk factors for AD) remained unknown. Here, in this manuscript, we investigate the effect of ACAT1/SOAT1 pharmaceutical inhibition *in vitro* APOE4 cells and *in vivo* aging APOE4 mice. We reported our finding in detail below.

## 2. Results

### 2.1 In vitro F12511 treatment in primary APOE4 microglia reduces lipid droplets and increase ABCA1 protein content

Since APOE4 expressing microglia are reported to have neutral lipid droplets accumulation and secrete higher level of proinflammatory cytokines, we first chose to study the effect of ACAT1/SOAT1 pharmaceutical inhibition in APOE4 primary microglia [10,11]. Previously, ACAT1/SOAT1 inhibitors were demonstrated to reduce lipid droplets and produce anti-inflammatory response in both microglia and macrophages, which makes them an attractive candidate for intervention in APOE4 microglia [26,29-31]. To establish an in vitro system for studying the effect of ACAT1/SOAT1 inhibition, we first isolated primary microglia from APOE3-TR and APOE4-TR mice following procedure described in Figure 1A. We then exposed these microglia to ACAT inhibitor F12511 as shown in Figure 1B. We chose F12511 for this study, as our laboratory had previously developed a drug delivery system using F12511 for ACAT1/SOAT1 inhibition in vivo [28]. Primary microglia were allowed to rest in serum free media overnight and all treatment was performed in serum free condition to ensure primary microglia transcriptome is more native microglia like [32].

**Figure 1.**
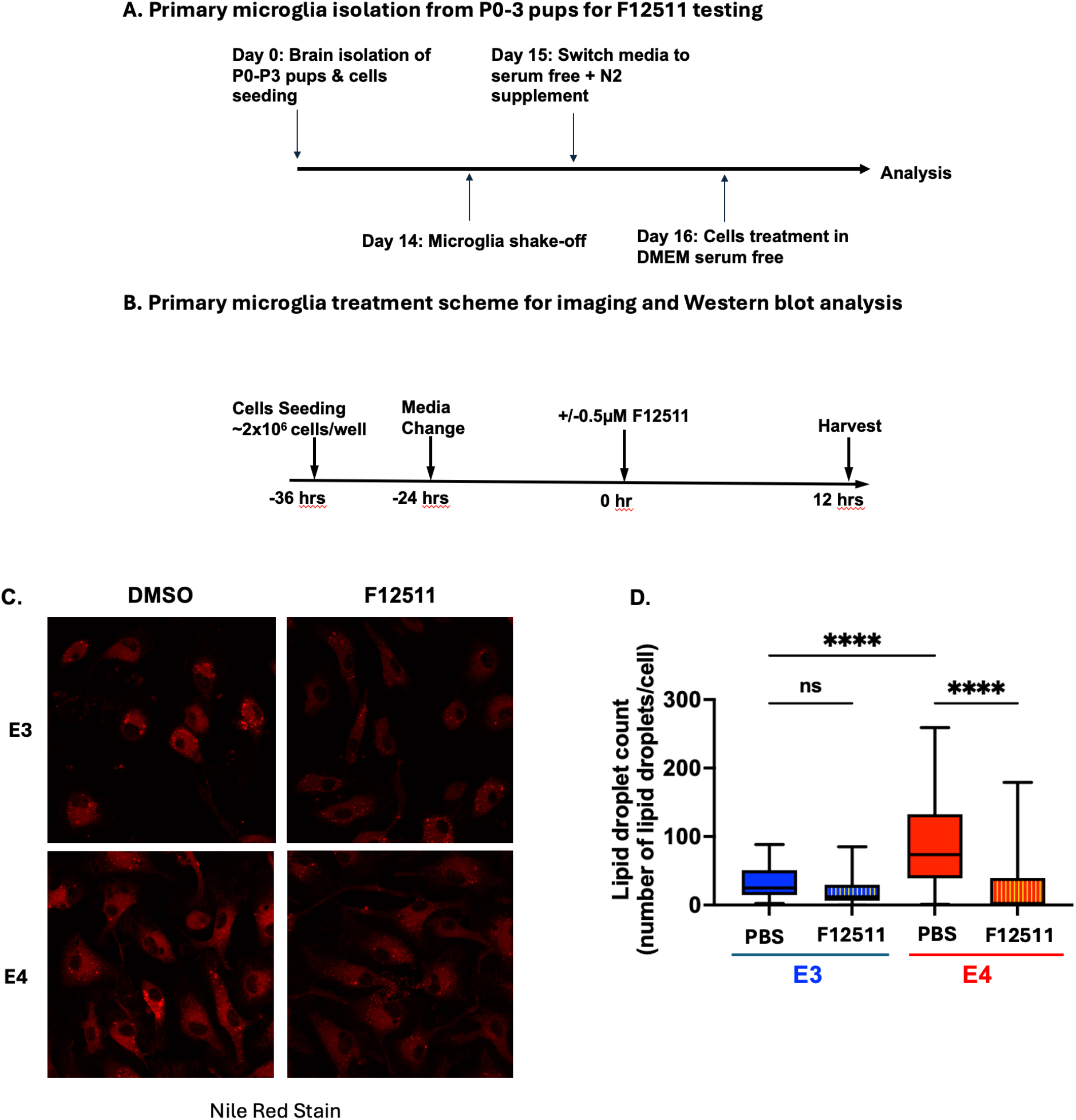
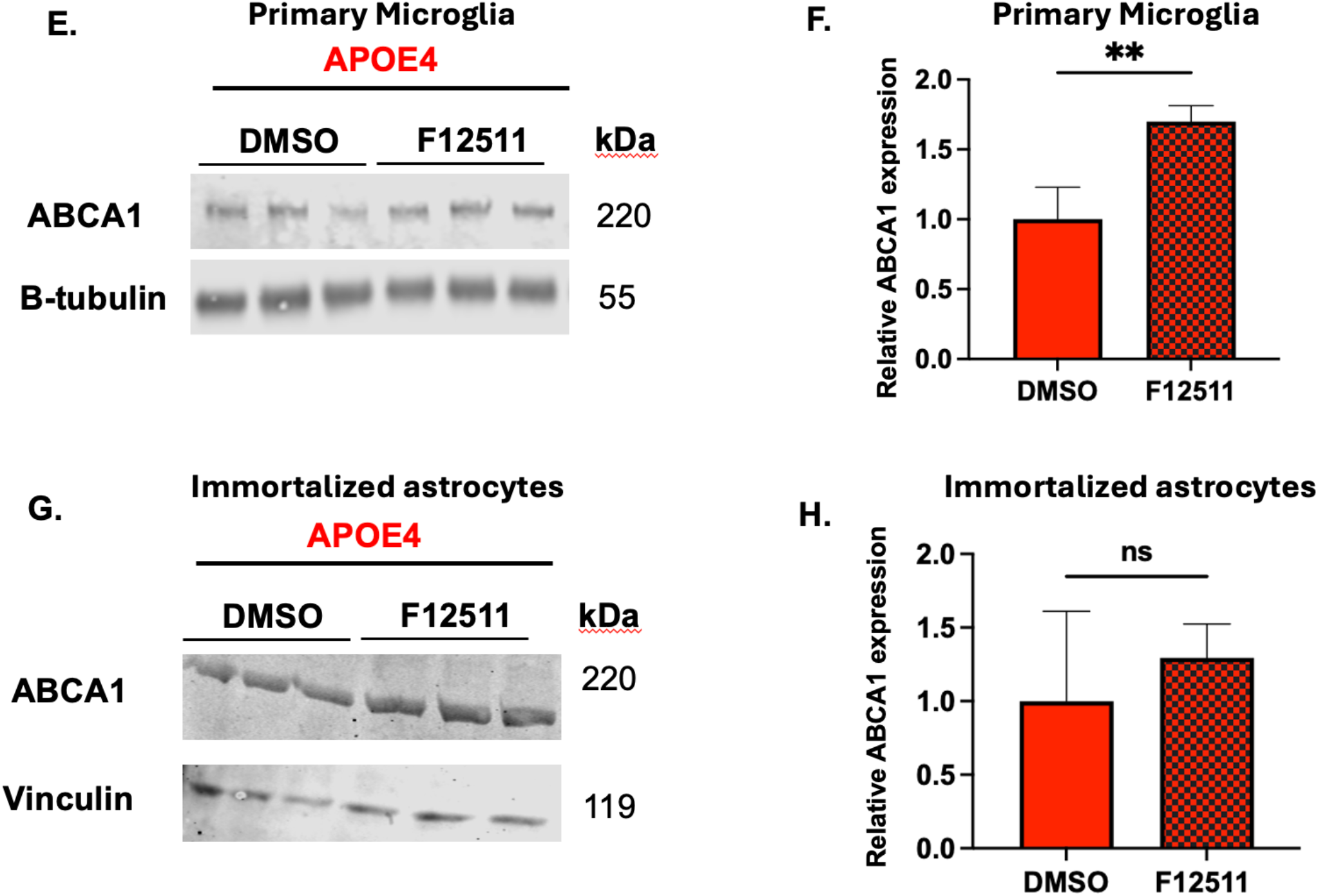
*In vitro* F12511 treatment reduces lipid droplets and upregulates ABCA1 protein content in primary APOE4 microglia, but not in APOE4 immortalized astrocytes. (A) Experimental time-line for primary microglia isolation from P0-3 pups. (B) Primary microglia treatment scheme for imaging and western blot analysis in serum free media. (C) Nile Red assay in primary APOE3 and APOE4 microglia treated with or without F12511. (D) Lipid droplets quantification from Nile Red data. N = 40 cells per treatment group were analyzed. (F) Representative western blot monitoring ABCA1 protein content in APOE4 primary microglia treated with or without F12511. (E) Quantification of western blot data. (G) Representative western blot monitoring ABCA1 protein content in APOE4 immortalized astrocytes treated with or without F12511. (H) Quantification of western blot data. N =3 for western blot experiments. Value of cells treated with DMSO was normalized to 1. Data expressed as mean +/- SEM. **P < 0.01; ****P < 0.0001; ns: not significant.

We first tested whether ACAT1/SOAT1 inhibition would reduce lipid droplets in APOE4 primary microglia by treating cells with F12511 for 12 hrs and staining them with Nile Red. Nile Red is a dye that is widely used for visualizing intracellular lipid droplets (contain mainly CE and TAG) [33]. Nile Red can be used for monitoring the effect of ACAT1/SOAT1 inhibition [34,35]. We reported Nile Red stained cells representative images and quantification results in **Figure 1C and 1D**. Our result showed that APOE4 microglia exhibited a higher lipid droplets count compared to APOE3 primary microglia, suggesting APOE4 primary microglia has more neutral lipid droplets at baseline. This result agrees with studies from Machlovi et. al and Litvinchuk et. al [10,11]. Treatment with ACAT1/SOAT1 inhibitor F12511 did not significantly reduce lipid droplets count in APOE3 primary microglia, as their lipid droplets level is already low. While in APOE4 primary microglia, F12511 treatment significantly reduced lipid droplets count. These results indicated that besides TAG as previously reported in literature, there is also higher level of CE in APOE4 primary microglia compared to its APOE3 counterpart and this CE rich lipid droplets pool in APOE4 primary microglia is sensitive to ACAT1/SOAT1 inhibition.

We next asked what the consequence of CE lipid droplets reduction in APOE4 primary microglia is when ACAT1/SOAT1 is inhibited. Previously, we demonstrated that in human microglia HMC3 cells loaded with cholesterol rich myelin debris, ACAT1/SOAT1 inhibition reduced CE accumulation, and increased cholesterol efflux via upregulation of ATP transporter cassettes A1 (ABCA1) gene expression and protein content [29]. Since APOE4 primary microglia was reported to have disrupted cholesterol metabolism and increased lipid droplets accumulation, we speculated that ACAT1 inhibition might also upregulate ABCA1 content in APOE4 primary microglia [8,10,11]. (To test, we treated the APOE4 primary microglia with F12511 for 12 hours, followed by monitoring ABCA1 protein content on Western blot and quantified ABCA1 signal. **(Figure 1E and 1F)** Our result reveals that ABCA1 protein content increased with F12511 treatment, suggesting ACAT1 inhibition might also alleviate cholesterol burden in APOE4 primary microglia cells through activating ABCA1 transporter [29].

As different cell types in the CNS play different roles in cholesterol trafficking and metabolism, it is possible that the effect of F12511 is cell-type dependent. Astrocytes are responsible for cholesterol synthesis in the CNS and in APOE4 expressing cells, cholesterol accumulates in the lysosome [8]. To test whether F12511 can also increase ABCA1 protein in astrocytes, we treated immortalized APOE4 astrocytes with F12511 and monitor ABCA1 protein content on western blot **(Figure 1G, H)**. Unlike in microglia, our result showed that F12511 treatment in APOE4 astrocytes did not impact ABCA1 protein content. In APOE4 immortalized astrocytes, ABCA1 protein has been previously reported to be lower than its APOE3 counterpart [36]. APOE4 expression in astrocytes triggers ABCA1 aggregation in the late endosome and prevent its recycling to the plasma membrane in an ADP-ribosylation factor 6 (ARF6) dependent manner [36]. It is possible that blocking ACAT1 does not bypass ARF6 pathway to increase ABCA1 expression, unlike what we observed in microglia. Overall, our result suggests that the effect of F12511 is cell-type dependent.

### 2.2 Pharmaceutical inhibition of ACAT1 by F12511 dampen NFκB activation in primary E4 microglia in a TLR4 dependent manner

Next, to understand the consequence of CE rich lipid droplet reduction and ABCA1 protein upregulation through ACAT inhibition, we evaluate APOE4 microglia response to inflammation stimulus using lipopolysaccharide (LPS). LPS is considered a pan -inflammation agent; it is a TLR4 specific ligand in microglia and is commonly used for studying TLR4 and its signaling pathway.

ACAT inhibition and ABCA1 protein upregulation can alter membrane cholesterol content. Cholesterol regulates immune signaling through modulating lipid rafts (Reviewed in [37]). Lipid rafts are microdomains that are cholesterol rich and serve as platforms to carry out different receptors signaling events. Altering cholesterol content within lipid raft microdomain regulate membrane’s fluidity and immune receptors signaling such as TLR4 (Reviewed in [37]). We had previously shown that ACAT1/SOAT1 inhibition in LPS-treated N9 cell promotes TLR4 endocytosis from the plasma membrane to the endolysosomal compartment for degradation, likely by regulating cholesterol content in TLR4 lipid raft microdomain [26]. Additionally, the intracellular juxtamembrane of TLR4 was shown to contains a cholesterol recognition amino-acid consensus sequence, suggesting the role of cholesterol directly in TLR4 regulation (Reviewed in [37]). Evidence suggested that APOE4 might carry out damaging phenotypes in AD in a TLR4-dependent pathway [38-40]. LPS brain infusion in APOE4 mice results in increasing NFkB activation and il-1β secretion compared to APOE3 counterpart [39,40].

In LPS induced N9 mouse microglia, ACAT1/SOAT1 pharmaceutical inhibition by using a different ACAT1/SOAT1 inhibitor K604 dampen NFκB activation and proinflammatory via modulating the fate of TLR4 [26]. Thus, we speculate that ACAT1/SOAT1 inhibition in APOE4 microglia might also be beneficial by dampen proinflammatory response in a TLR4 dependent manner. To test this possibility, we monitored NFκB activation in LPS treated APOE4 microglia with or without F12511 treatment. Cells without LPS exposure were used as control. To further validate whether F12511 effect is TLR4 dependent, we also treated cells that are exposed to LPS with or without F12511, with a TLR4 specific inhibitor – TAK242 [41]. Treatment scheme and experimental set up is graphically illustrated in **Figure 2A**. Phosphorylation state of p65 and iκβ-*α* were measured to evaluate NFκB activation state [42]. Representative western blot is shown in **Figure 2B** and quantification of these two markers are reported in **Figure 2C** (p65) and **Figure 2D** (iκβ-*α*). Our analysis revealed that F12511 treatment in APOE4 microglia dampens NFκB activation when looking at both markers. **(Third and fourth bar, Figure 2C and 2D**) Treatment of these cells with TAK242 abolished F12511 effect in APOE4 cells exposed with LPS, suggesting F12511 effect on dampening NFκB activation is dependent on TLR4 signaling. **(Fifth and sixth bar, Figure 2C and 2D)** Overall, our result demonstrated that F12511 treatment in APOE4 primary microglia cells dampen NFκB activation in a TLR4 dependent manner. This result agrees with our previous published data in N9 cells carrying mouse APOE and using a different ACAT inhibitor K604 [26]. Together, both of these data suggest that ACAT blockade dampen NFκB through TLR4 pathway is conserved regardless of APOE isoform was from human and mouse.

**Figure 2.**
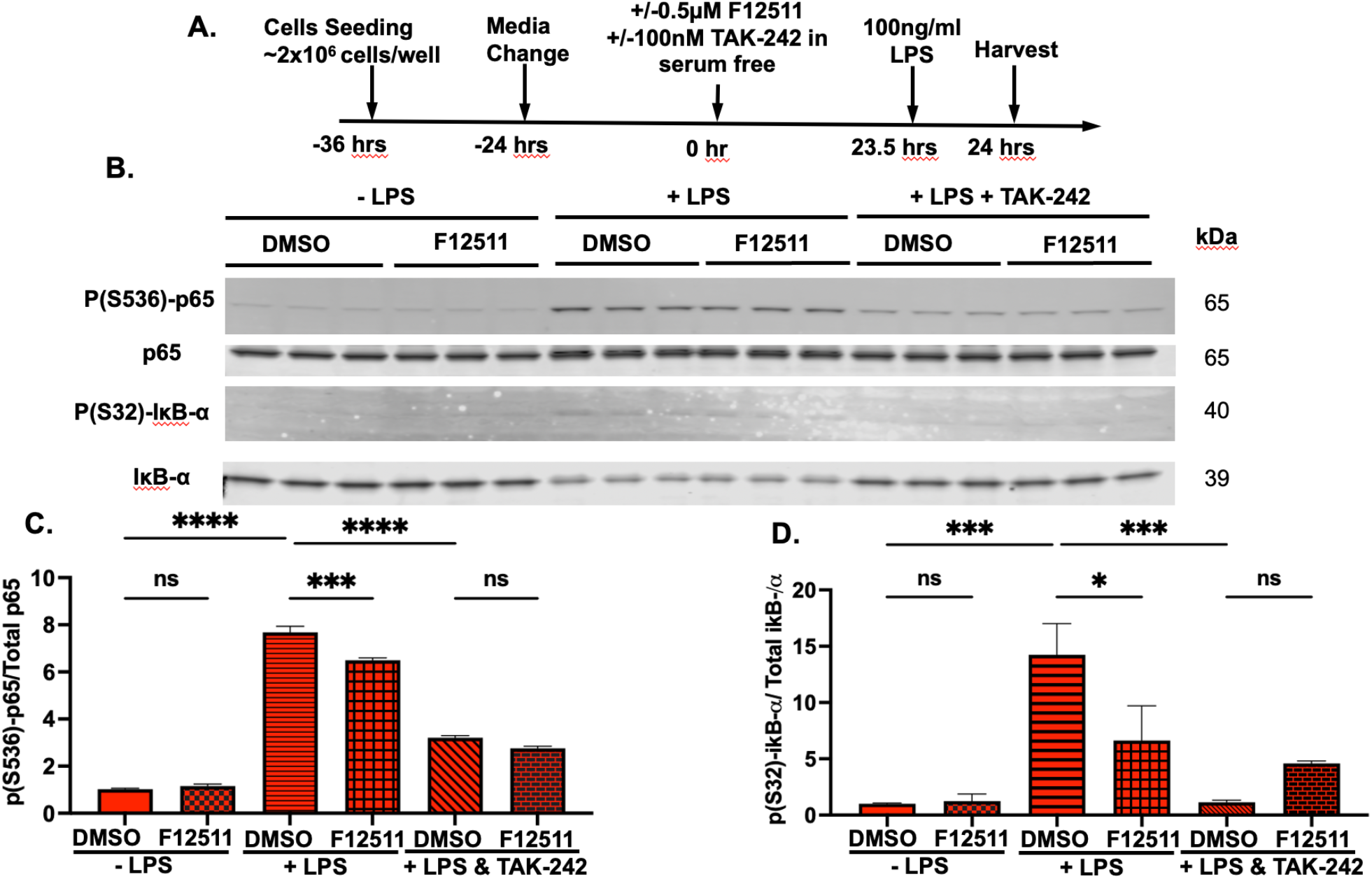
Pharmaceutical inhibition of ACAT1 by F12511 dampen NFκB activation in LPS induced primary E4 microglia in a TLR4 dependent (TAK-242 sensitive) manner. (A) Treatment scheme. (B) Representative western blot of keys NFκB activation markers P(S536)-p65, p65, P(S32)-IκB-*α*, IκB-*α*. Western blot quantification for (C) P(S536)-p65/p65 ratio and (D) P(S32)-IκB-*α*/IκB-*α* ratio. N = 3. Value of cells treated with DMSO with no LPS was normalized to 1. Data expressed as mean +/- SEM. *P < 0.05, ***P < 0.001, ****P < 0.0001.

### 2.3 Design of F12511 in vivo efficacy studies in APOE3 and APOE4 at different age range

To test ACAT1/SOAT1 inhibitor efficacy in vivo, we employed the use of a lipid-based nanoparticle system encapsulating ACAT1/SOAT1 inhibitor F12511 that was previously published and tested in wild type (WT) and triple transgenic (3xTg) AD mice [28,43]. A graphical diagram describing this system is illustrated in **Figure 3A**.

**Figure 3.**
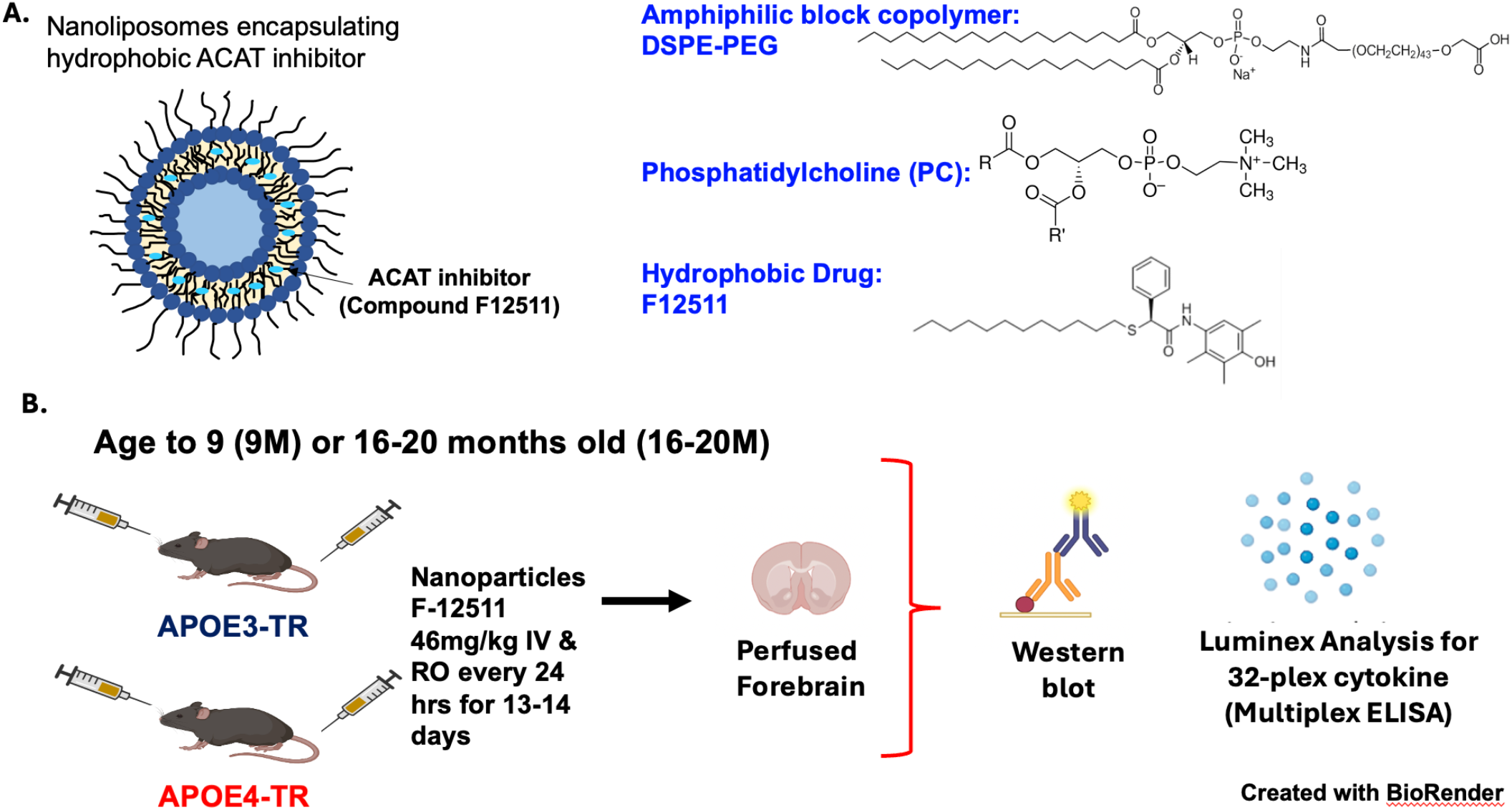
Design of F12511 in vivo efficacy studies in APOE3 and APOE4 at different age range. (A) Diagram and components of nanoparticle encapsulating ACAT1/SOAT1 inhibitor. Nanoparticles are made up of DSPE-PEG and PC and ACAT1/SOAT1 inhibitor F12511 according to procedure published in De La Torre et. al 2021 (PMID: 34890698). (B) Animal treatment scheme. Mice were aged to 9 months old (9M) or 16-20 months old (16-20M) for this study, followed with daily injection by alternate intravenous (IV) and retro-orbital (RO) route. Mice were then perfused with PBS, and forebrain tissues were collected and homogenized for western blot and Luminex analysis.

Employing the use of our nanoparticle system, we then injected adult (9M old) and aged (16-20M old) female APOE3-TR and APOE4-TR mice with PBS (control), nanoparticle (vehicle) and nanoparticle F12511 daily for 13-14 days. We chose to use exclusively female mice in this study, as other studies reported that female APOE4 mice tend to experience worsen neuroinflammation compared to its male counterparts [44,45]. We collected forebrain tissues and analyzed key inflammatory markers using Luminex xMAP technology and Western blot. As AD is an aging related disease, we designed our studies to examine the effect of nanoparticle F12511 in APOE4-TR adult and aged mice to determine nanoparticle F12511 efficacy at two different disease stage.

### 2.4 Two weeks daily IV injection of nanoparticle F12511 at 46mg/kg subtly changes inflammatory profile in the forebrains of 9M old APOE4 mice, but not APOE3 mice

We first examined the effect of nanoparticle F12511 in 9M old adult APOE3 and APOE4 mice. Since we saw acute 12 hrs treatment with F12511 was able to dampen NFκB activation in a TLR4 dependent manner **(Figure 2)** and treatment with another ACAT1/SOAT1 inhibitor K604 was reported to reduce TLR4 protein content in N9 microglia after 48h treatment [26], we speculate that with two weeks daily treatment, nanoparticle F12511 might decrease TLR4 protein content in the brain.

After treatment with nanoparticle F12511, we observed a slight decrease in TLR4 protein content in 9M old APOE4 mice forebrain, but not in their APOE3 counterpart **(Figure 4A)**. To determine whether this drop in TLR4 protein content in APOE4 mice correlate with proinflammatory response in the forebrain region, we analyzed the same tissues using Miliplex xMAP technology to assess different pro and anti-inflammatory cytokines level. The analyzed cytokines within detectable range are reported in **Figure 4B**. Proinflammatory cytokines such as il-1*α*, TNF*α*, il12p70 that are downstream of NFκB activation did not change significantly. Interestingly, nanoparticle F12511 treatment in APOE4 mice slightly increases cytokines such as il-9 and il-13 compared to PBS treated mice. The role of il-9 and il-13 in APOE4 mice is unknown, however, il-13 is generally recognized as an anti-inflammatory cytokine. In an amyloid precursor protein (APP) mouse model for AD – APP23, intracerebral injection of il-4/il-13 increases Aβ clearance and improves cognitive deficits [46]. Additionally, it is unknown how il-13 and TLR4 signaling correlates in microglia, but in epithelial cells, il-13 decreases TLR4 function [47].

**Figure 4.**
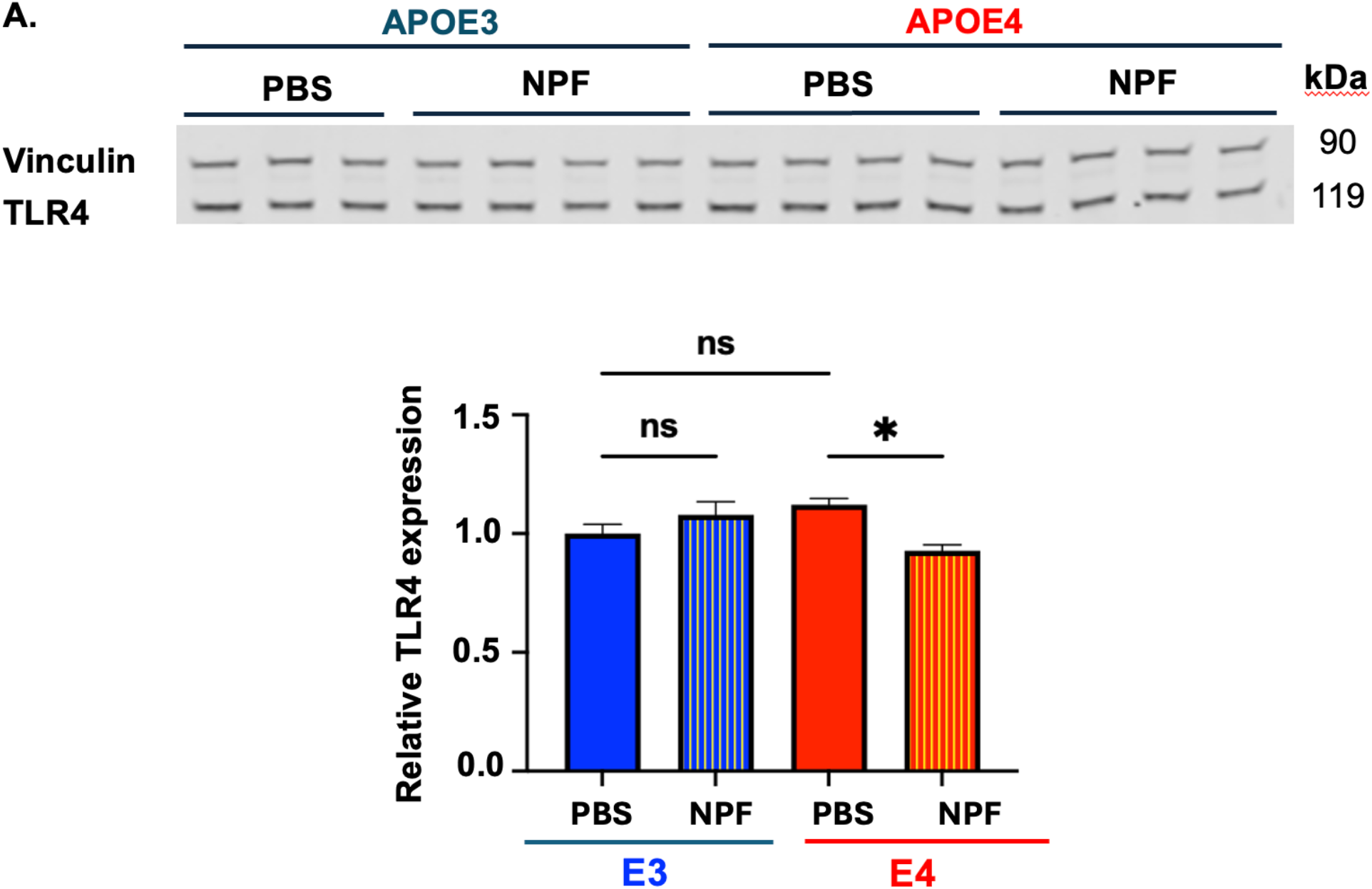
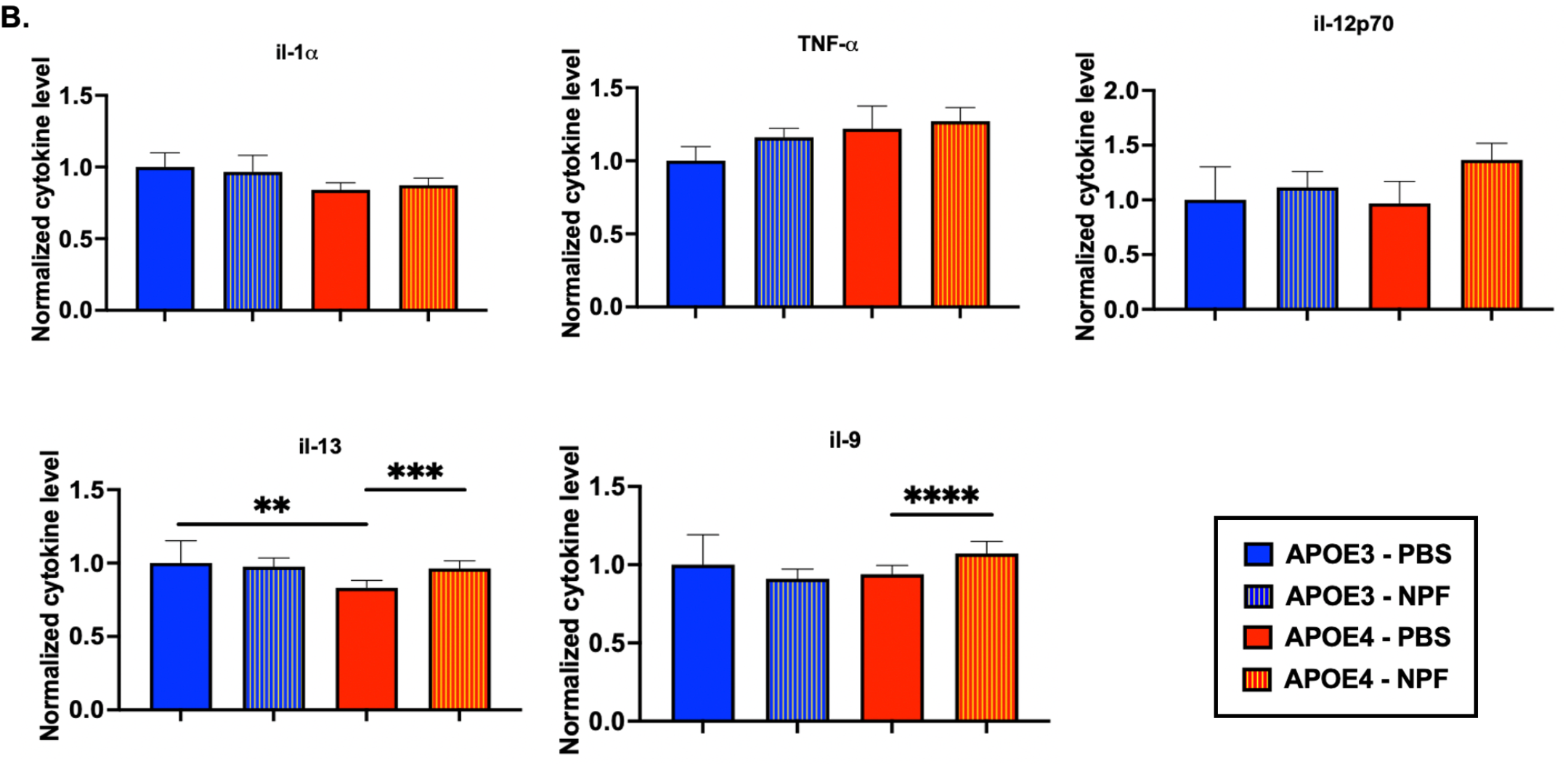
Two weeks daily alternate IV and RO injection of nanoparticle F12511 at 46mg/kg subtly changes in 9M old APOE4 mice inflammatory profile, but not in APOE3 mice. (A) Representative western blot and quantitative analysis of relative TLR4 expression in 9M old injected forebrain homogenate. Quantitative analysis revealed nanoparticle F12511 slightly reduced total TLR4 protein expression in APOE4 forebrain homogenate, but not in APOE3 mice. (B) Detectable Alzheimer’s related cytokines and cytokines that are significantly altered by drug treatment from MILLIPLEX xMAP analysis were plotted individually. IL-13 and IL-9 changed significantly in APOE4 treated forebrain homogenates, while APOE3 mice were unaffected. Value of APOE3 mice treated with PBS was normalized to 1. Data expressed as mean +/- SEM. **P < 0.01,***P < 0.001, ****P < 0.0001.

In adult mice, we observed subtle effect from nanoparticle F12511 treatment in APOE4 mice but not APOE3 mice. This difference might be due to APOE4 mice exhibiting a leaky BBB as part of its disease phenotype [48], which allows more nanoparticle to enter the brain cells and alter the brain’s inflammatory profile. We next examine the effect of nanoparticle F12511 treatment in aging APOE3 and APOE4 mice, where the BBB is leaky as aging progresses.

### 2.5 Two weeks daily alternate IV injection of nanoparticle F12511 at 46mg/kg changes PLIN2, TLR4 protein content and inflammatory profile in the forebrain in 16-20M old APOE3 and APOE4 mice

As APOE4 phenotype exacerbates by age, we aged APOE3 and APOE4 mice to 16-20M old to test the effect of nanoparticle F12511. We previously saw in 3xTg AD mice, nanoparticle F12511 carrying out prominent effects in 16-20M old age group [28].

To test whether nanoparticle F12511 treatment reduces neutral lipids droplet in vivo, we monitored perilipin-2 (PLIN2) protein content in total forebrain lysate. **(Figure 5A)**. Previous, it was shown that a decrease in PLIN2 signal is associated with decreasing in lipid droplet content [10,49]. As expected, similar to our data in APOE4 primary microglia, we observed that PLIN2 protein content in the forebrain of nanoparticle F12511 treated group exhibit a decreasing trend compared to the PBS treated group in both APOE3 and APOE4 mice, suggesting a reduction in neutral lipid droplet content in the brain. This result agrees with previous published data by Valencia-Olvera et. al, using a different ACAT inhibitor (avasimibe) and administered through a different route in APOE4/5xFAD mice [27]. In principle, reduction in brain CE rich lipid droplets release free cholesterol that can participate in important cellular process [50].

**Figure 5.**
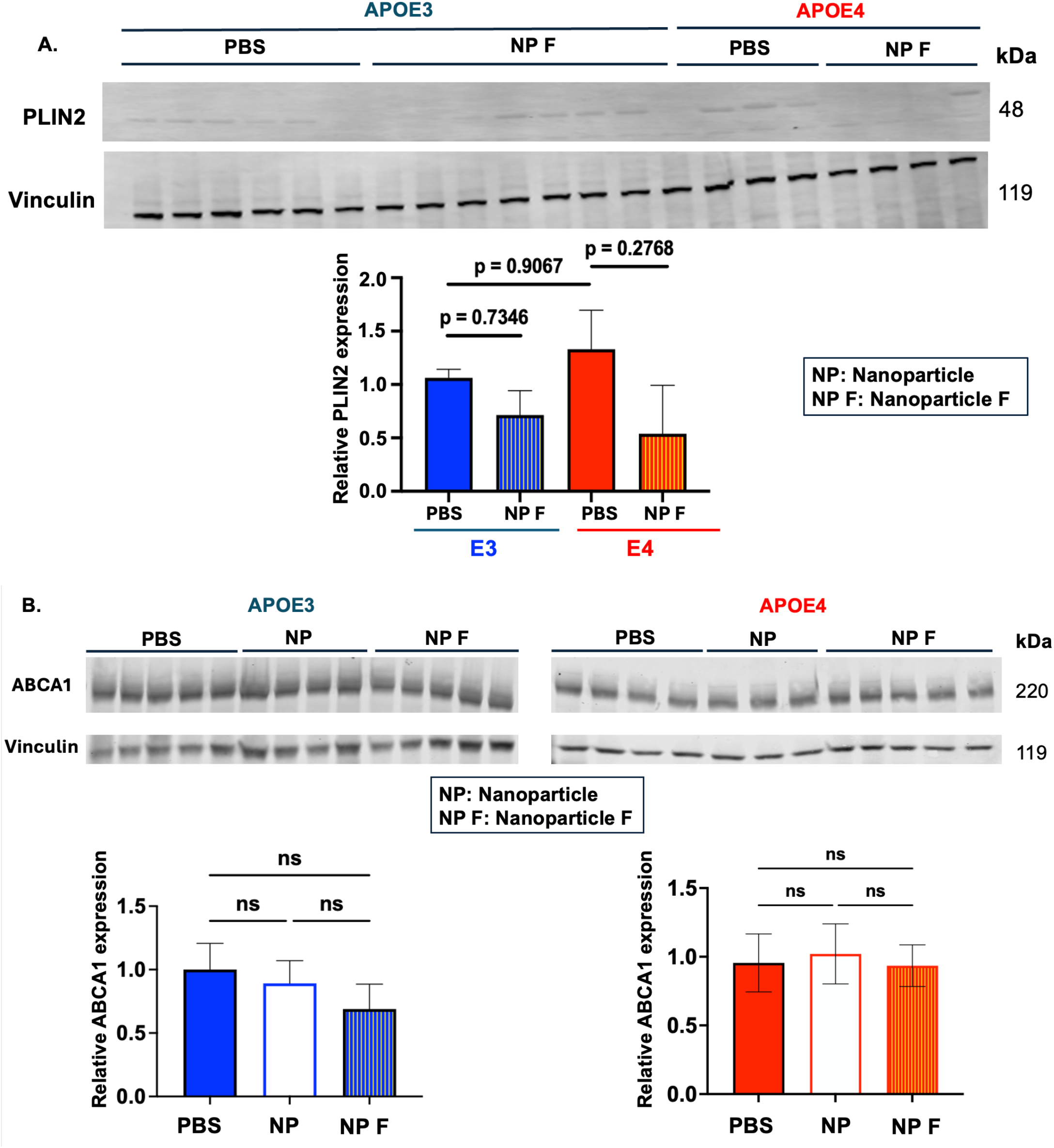
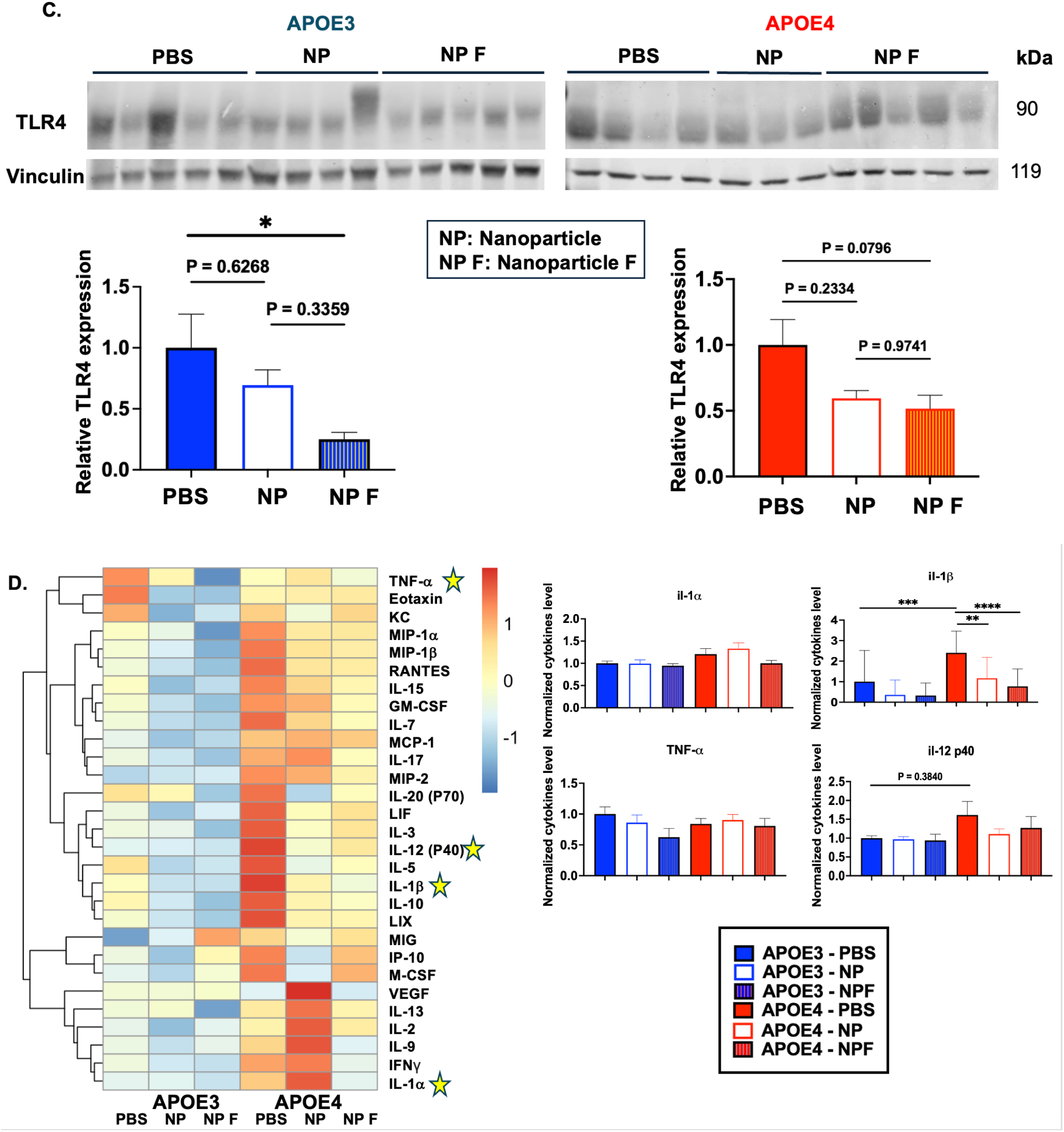
Two weeks daily alternate IV and RO injection of nanoparticle F12511 at 46mg/kg significantly alter inflammatory profile and lipid droplet markers in both APOE3 and APOE4 forebrain at 16-20M old. (A) Representative western blot and quantitative analysis of PLIN2 protein expression. (B) Representative western blot and quantitative analysis of ABCA1 protein expression (C) Representative western blot and quantitative analysis of relative TLR4 protein expression in 16-20M old injected forebrain homogenate. (D) Heatmap visualizing average cytokines readings from each treatment groups with Z-score transformation. (left panel) Detectable Alzheimer’s related cytokines and cytokines that are significantly altered by drug treatment from MILLIPLEX xMAP analysis were plotted individually (right panels). Value of APOE3 mice treated with PBS was normalized to 1. Data expressed as mean +/- SEM. *P < 0.05, **P < 0.01,***P < 0.001, ****P < 0.0001.

We also determined the effect of nanoparticle F12511 on ABCA1 protein content in vivo **(Figure 5B)**. Our result demonstrated that in aging APOE3 and APOE4 mice, both nanoparticle and nanoparticle F12511 did not affect ABCA1 protein content. This data is in agreement with our study in cell culture. in APOE4 microglia and astrocytes **(Figure 1 E,F,G,H)**. Together, these results suggest the effect of F12511 on ABCA1 protein is cell-type dependent. While F12511 increases ABCA1 protein content in APOE4 microglia, it might not be able to induce ABCA1 protein expression in APOE4 astrocytes, which counts for the majority of cells in the brain.

We observed a significant decrease in TLR4 protein content in aging APOE3 mice and a trending decrease in TLR4 protein content in aging APOE4 mice. **(Figure 5C)** This finding agrees with previous data from Li et. al [26]. Li et al showed that in LPS induced N9 microglial cells, acute treatment (∼4 hours) with the ACAT1 inhibitor K604 modulates the fate of TLR4 and increases TLR4 endocytosis rate to the lysosome; while chronic treatment of ACAT inhibitor (48 hours) results in TLR4 protein degradation in the lysosomes [26]. Our result suggested that 2 weeks daily injection is sufficient to decrease TLR4 protein content in both aging APOE3 and APOE4 forebrain.

Finally, to determine whether the reduction of TLR4 protein content and decrease in neutral lipid droplets correlate with inflammatory profile in the brains of E3/E4 mice, we analyzed aged 16-20M old forebrain tissue cytokines with Milliplex xMAP technology, in similar manners with 9M old mice samples. Results from this screen is reported in **Figure 5D**: **left panel** compare average cytokine level between different treatment groups. As expected, 26 out of 31 screened cytokines are elevated in APOE4 mice compared to APOE3 mice, which suggested a more proinflammatory phenotype in APOE4. This result is consistent with previous literature (first and fourth column from the left, **Figure 5D, left panel** [10,11,51]. Treatment with nanoparticle F12511 results in a decreasing trend in the majority of proinflammatory cytokines in both APOE3 and APOE4 mice compared to PBS and nanoparticle alone treated groups **(Figure 5D, left panel)**. In some cytokines, treatment with nanoparticle alone decreases cytokines level compared to PBS and further decreases with nanoparticle F12511 such as TNF-α in APOE3 and il-1β in APOE3 and APOE4, suggesting that there might be additive effect of nanoparticle and F12511 **(Figure 5D, right panel)**. We plotted individual cytokines that are related to AD and neuroinflammation such as il-1α, il-1β, TNF-α, il-12p40 to further assess their inflammatory profile, and observed decreasing trends of these cytokines in APOE4 mice treated with nanoparticle F12511 compared to PBS treated or nanoparticle treated mice **(Figure 5D, right panel)**. Specifically, the result shows that out of all cytokines, nanoparticle F12511 treatment has the most significant effect on decreasing il-1β in APOE4 mice. This observation is consistent with our previous data in Li et. al and **Figure 5C** [26]. Since il-1β is released down-stream of TLR4 activation, therefore decreasing in TLR4 content is expected to result in lower level of il-1β secretion. Together, our data in **Figure 5** support the conclusion that nanoparticle F12511 decreases TLR4 protein content, lipid droplets and reduce proinflammatory cytokines in both aged APOE3 and APOE4 mice brains.

When inhibiting ACAT1 in vivo with nanoparticle F12511, we observed a more pronounce effect in aging mice compared to adult mice **(Figure 4B and 5D)**. This result could likely be due to two possibilities: (1) The BBB is leaky in aging mice, which allows more drug to enter the brain to carry out its therapeutic effect, (2) APOE4’s pathological phenotype is more significant with age, which leads to more drastic difference in treated group compared to control group. Previously, we demonstrated that in 3xTg mice, nanoparticle F12511 treatment also result in a more notable therapeutic effect in aging mice compared to adult mice, which is consistent with what we observed in APOE4 mice here [28]. Additionally, our data in **Figure 5D, left panel** showed that in aging mice at 16-20M old of age, APOE4 mice exhibits more inflammation in the brain compared to its APOE3 counterparts, where this difference is more subtle in adult mice at 9M old of age. This result is consistent with previous reported data [51].

Interestingly, we noticed that in aging mice, nanoparticle with PC but without F12511 exhibited suppressive effect on TLR4 and il-1β. **(Figure 5A and Figure 5C, right panel)** We had previously made a similar observation in aging (16-20 months of age) 3xTg mice: nanoparticle with PC but without F12511 had a reduction effect on human APP and human hyperphosphorylated Tau, and a reduction effect on proinflammatory cytokines level in selective cytokine markers, such as Eotaxin and TNF-*α* [28]. These observations together suggest that nanoparticle with PC and F12511 may work in additive manner to suppress neuroinflammation. At present, it is unclear which cell type(s) are responsible for the effect by nanoparticle with PC without F12511. We speculate that one of the primary targets for nanoparticle with PC could be vascular endothelial cells, as they are exposed directly to nanoparticle with PC upon their injection into the bloodstream. Once the BBB becomes leaky, nanoparticle will then be exposed to other cell types like pericytes, astrocytes, microglia, neurons and oligodendrocytes. Phosphatidylcholine (PC), a component of our nanoparticle system, was demonstrated to exhibit anti-inflammatory properties in LPS-induced systemic inflammation model [52]. It is possible that nanoparticle anti-inflammatory response observed in this study is due to PC effect. Additionally, PC can alter lipid raft size, which serves as a signaling platform for TLR4 [53]. Changes in lipid raft can regulate receptors-mediated endocytosis and degradation of different proteins, which could also explain our TLR4 result. To elucidate this hypothetic mechanism, future study focuses on understanding the effect of nanoparticle formulated with and without PC is necessary.

## 3. Discussion

In this study, we aimed to investigate the effect of ACAT1 inhibition in APOE4 cell and mouse models. We showed that in vitro, ACAT1 inhibition in APOE4 primary microglia decreases CE rich lipid droplets, upregulates ABCA1 protein content and dampen NFkB mediated activation in a TLR4 dependent manner **(Figure 1 & 2)**. ACAT1 inhibition does not impact ABCA1 protein content in APOE4 astrocytic cell line **(Figure 1)**. We then tested the effect of inhibited ACAT1 in vivo by using the lipid nanoparticle system containing DSPE-PEG_2000_, PC and ACAT1 inhibitor F12511 on female adult (9M of age) and aging (16-20M of age) mice **(Figure 3)**. Our results show that nanoparticle F12511 decreases TLR4 protein content, decreased lipid droplets and dampen proinflammatory cytokines secretion in aging mice, while the effects nanoparticle F12511 in same set of mice at adult age is more subtle **(Figure 4 & 5)**.

In the brain, it is believed that APOE is involved in intracellular membrane cholesterol trafficking and recycling, in a cell type specific manner. APOE4 is defective in intra-cellular cholesterol trafficking, causing inefficient utilization of free cholesterol, which leads to cholesterol accumulation in the ER, and subsequently leads to CE formation in cells [8,9,11]. Interestingly, a general mechanism has been proposed to account for the action of APOE4: In various cell types, APOE4 alters the pH of endosome and disrupts endosomal membrane recycling function [54].

CE and TAG are the two main components of lipid droplets. While other studies emphasized on lipid droplets accumulation in APOE4 cells, specifically TAG accumulation. To our knowledge, studies focusing on the importance of removing CE droplets and how that may benefit APOE4 carriers have been rare. Haney et. al showed that inhibiting ACSL1, a protein that is upstream of ACAT1 (responsible for CE synthesis) and DGAT (responsible for TAG synthesis) in APOE4 mice reduce disease pathology and ameliorate neuroinflammation [12]. ACSL1 is also responsible for the bulk synthesis of phospholipids which is a main ingredient of all cell membranes. The ACSL1 inhibitor safety profile is a concern, which poses a need for an alternative approach to tackle lipid droplets accumulation in APOE4 model [55]. Another study by Blanchard et. al removed CE lipid droplets in APOE4 mice by using cyclodextrin injection, cyclodextrin is a non-specific cholesterol sequester that helps to remove cellular cholesterol [9]. High concentration of cyclodextrin treatment tend to cause rodents to become deaf. Our current work shows that removing CE is beneficial in the APOE4 model, and this can be accomplished by specifically inhibiting ACAT1 using F12511, its safety was approved by FDA. Valencia-Olvera et. al used another FDA safely approved ACAT inhibitor CI-1011 in APOE4/5xFAD model, and showed CI-1011 dampen neuroinflammation in these mice [27]. However, their work cannot assure that ACAT1 inhibition still benefit APOE4 disease phenotype when the subjects are without the AD background (5xFAD). Our finding described here filled in the gap of knowledge, demonstrating that in the absent of Aβ, ACAT1 inhibition is still beneficial in APOE4 mice in terms of ability to suppress neuroinflammation. We used APOE4 primary microglia as a cell model to study the mechanism of ACAT1 inhibition and showed that the effect of ACAT1 inhibition link with two membrane receptors at the PM: TLR4 and ABCA1. This mechanism may also be applied to other cell types in the brain, including cells at the blood brain barrier: (i.e., vascular endothelial cells, pericytes, etc.) This possibility remains an area of investigation. These studies need to focus on how ACAT1 inhibition affect APOE4 phenotype in cell type specific manner in vivo. Overall, our study suggests that ACAT1 inhibition is a novel strategy to ameliorate CE rich droplets and dampen proinflammatory response in the aging APOE4 patients.

## 4. Materials and Methods

### 4.1 Animals

APOE3 and APOE4 target replacement (APOE3-TR and APOE4-TR) mice on a C57B6/J background were obtained from Taconic (La Jolla, CA, USA). Female mice aged 9 months and 16-20 months were used for in vivo studies. P0-3 pups (mixed gender) were used for primary microglia isolation. All mouse procedures were approved by the Dartmouth Institutional Animal Care and Use Committee.

For in vivo studies, female APOE3-TR and APOE4-TR mice within the indicated age range were treated daily for 13-14 days by alternating between intravenous injections via the tail vein and retro-orbital venous injections at a dose of 46 mg/kg nanoparticle F12511. Mice were then perfused with 30 ml of ice-cold PBS (MilliporeSigma, St. Louis, MO, USA) to remove blood contaminants. Forebrain tissues were collected and snap-frozen on dry ice. Tissues were stored at -80°C until further processing.

### 4.2 Cell culture

Primary microglia were isolated from P0-P3 pups according to procedures previously described in [56]. Briefly, brains were isolated, and meninges were removed. The brains were homogenized by mechanical pipetting and trypsinization, then plated on poly-L-lysine-coated flasks. Cells were cultured in DMEM with 10% heat-inactivated fetal bovine serum (FBS) and 1% penicillin/streptomycin (P/S), all from Gibco (ThermoFisher, Waltham, MA, USA), overnight. The next day, cells were rinsed with PBS and cultured in DMEM supplemented with 10% FBS, 1% P/S, and 1% L929 conditioned medium (growth factors) for 10 to 14 days. Primary microglia were harvested by shaking at 225 rpm at 37°C for 30 minutes and plated overnight in DMEM supplemented with 10% FBS and 1% P/S. Prior to treatment, cells were switched to DMEM free of serum and antibiotics. Immortalized mouse astrocytes expressing human APOE3 and APOE4 were a gift from Dr. David Holtzman, grown in DMEM/F12 supplemented with 10% FBS, 1% P/S (ThermoFisher, Waltham, MA, USA) All cells were maintained at 37°C with 5% CO_2_ in a humidified incubator.

### 4.4. Nile Red staining in live cells and images analysis

Nile Red was used to monitor lipid droplets in live cells according to procedures previously described in [35]. Briefly, primary microglia were plated on MaTek (Ashland, MA, USA) 35 mm dishes pre-coated with poly-L-lysine at 2×10^6^ cells per plate overnight. Treatment was performed in serum-free DMEM, and cells were rinsed three times with HBSS (Gibco by ThermoFisher, Waltham, MA, USA). Cells were treated with 100 ng/ml Nile Red and incubated for 10 minutes at 37°C, 5% CO_2_, protected from light. Cells were rinsed in HBSS and imaged in DMEM serum-free, with no phenol red (Gibco by ThermoFisher, Waltham, MA, USA), on the confocal fluorescence microscope. For image analysis, data were collected and analyzed using Fiji-ImageJ software version 2.1.0/1.53c, following the same pipeline described in [57].

### 4.5. Whole cell protein isolation, tissue homogenization and Western blot analyses

To obtain whole cell lysates, cells were harvested in RIPA buffer containing a protease inhibitor cocktail (MilliporeSigma, St. Louis, MO, USA) and incubated at 4°C for 30 minutes. For NFκB western blot, cells were lysed directly in sample buffers containing DTT, supplemented with protease inhibitor cocktails and PHOSStop phosphatase inhibitors (Roche, Indianapolis, IN, USA). RIPA cell lysates were then centrifuged, and the protein concentration of the supernatant was determined by Lowry protein assay.

To prepare brain homogenates, half of the frozen, perfused forebrains from female APOE3-TR and APOE4-TR mice within the indicated age ranges were homogenized with stainless steel beads in the Bullet Blender at 4°C in sucrose-based buffer, supplemented with a protease inhibitor cocktail as previously described in [25,28]. Brain homogenates were centrifuged at 12,000xg for 15 minutes, and supernatants were collected for western blot and Luminex analysis.

The lysates or homogenates were run on 6% SDS-PAGE gels or 4-20% Novex Tris-Glycine gels (ThermoFisher, Waltham, MA, USA) and transferred to a 0.45 μm nitrocellulose membrane for 4 hours at 300 mA, or a 0.2 μm PVDF membrane for 1 hour at 200 mA. Membranes were blocked in 5% milk in 1xTBS or 5% BSA in 1xTBS for phosphorylated protein detection. For primary antibody detection, the membranes were incubated overnight with anti-ABCA1 (Novus NB400-105), anti-PLIN2 (Proteintech 15294-1-AP), anti-TLR4 (Santa Cruz sc-293072), IκB*α* (CST, #4814), phospho-IκB*α* Ser32 (CST, 14D4), phospho-NFκB p65 Ser536 (CST, 93H1), NFκB p65 (CST, L8F6), and anti-Vinculin (Millipore 05-386) as a protein loading control. Blots were then washed and incubated with secondary antibodies from Li-Cor (Lincoln, NE, USA) with the appropriate species. Western blot images were captured on the Li-Cor Odyssey CLx and analyzed on Li-Cor Image Studio.

### 4.6. Preparation of nanoparticle and nanoparticle F12511

Nanoparticles F12511 were prepared according to procedures previously described in [28,43]. Briefly, DSPE-PEG2000 (Laysan Bio, Arab, AL, USA) in EtOH, L-*α*-Phosphatidylcholine (from egg yolk) (MilliporeSigma, St. Louis, MO, USA) in chloroform, and F12511 (Wuxi AppTec, Wuxi, China) in EtOH were combined, vigorously vortexed, and lyophilized overnight at -40°C, with the vacuum set at 133×10^−3^ mBar. Sterile PBS was then added to the lyophilized mixture so that the final concentration of the nanoparticle F12511 solution contained 30 mM DSPE-PEG2000, 6 mM PC, and 12 mM F12511. Mixtures were sonicated at 4°C for 2-4 cycles, with each cycle lasting 20 minutes. The sonicated solutions were spun at 12,000 rpm for 5 minutes, and supernatants were collected for animal injections. The resulting solution was protected from light throughout the entire process. For nanoparticles without the F12511 formulation, the solution was prepared following the same procedure, except F12511 was omitted.

### 4.7. Luminex analysis

Luminex analysis was performed using the procedure described previously in [28]. Cytokines from the mouse brain homogenates prepared in Section 4.5 were measured using Millipore mouse cytokine multiplex kits (EMD Millipore Corporation, Billerica, MA, USA). Calibration curves from the recombinant cytokine standards were prepared by following threefold dilution steps in the same matrix as the samples. High- and low-quality control samples with a known concentration provided by the manufacturer were used to validate the standard curve calculation. The standards and quality control samples were measured in triplicate, while the samples were measured once, and blank values were subtracted from all readings to ensure accurate measurement. All assays were carried out directly in a 96-well filtration plate (Millipore, Billerica, MA, USA) at room temperature and protected from light. Briefly, each well was pre-wet with 100 μL of PBS containing 1% BSA. Then, the beads, along with the standard, sample, quality control samples, or blanks, were added in a final volume of 100 μL. The plate was incubated at room temperature for 30 minutes with continuous shaking. The beads were washed three times with 100 μL of PBS containing 1% BSA and 0.05% Tween 20. A cocktail of biotinylated antibodies (50 μL/well) was added to the beads for a further 30 minutes of incubation with continuous shaking. The beads were washed three times and then streptavidin-PE was added for 10 minutes. The beads were again washed three times and resuspended in 125 μL of PBS containing 1% BSA and 0.05% Tween 20. The fluorescence intensity of the beads was measured using the Bio-Plex array reader 200 from Bio-Rad. Bio-Plex Manager software Version 6.2 with five-parameter curve fitting was used for data analysis.

### 4.8. F12511

was custom synthesized by WuXi AppTec in China, with verified stereospecificity and 98% purity.

### 4.9. Statistical analysis

All statistical analysis was performed using Prism10 software (GraphPad). A one way ANOVA test with Sidak multiple comparison was used to analyze data between treatment group. A two-way ANOVA with Turkey multiple comparison test was used for Luminex analysis. Error bars indicate SEM. * P<0.05; ** P<0.01; *** P<0.001; ****P<0.0001.

## Acknowledgements

We thank Dr. David Holtzman of Washington University at St. Lous for generously providing the human APOE3/E4 astrocyte cell lines. We thank Dr. Karl Biggs in Dr. Matthew Havrda’s lab for providing advice on primary microglia isolation. We thank Dr. Zdenek Svindrych for his guidance on microscope handling and imaging.

## Author contributions

T.N.H., C.C.Y.C. and T.-Y.C. designed research with input from M.H.; T.N.H., E.N.F performed research; T.N.H., analyzed data; T.-Y.C., C.C.Y.C., and T.N.H. wrote and edited the manuscript.

## Funding

This work was supported by NIH Grant R01 AG063544 (to T.-Y.C and C.C.Y.C.) and the Dartmouth PhD Innovation Program (to T.N.H). R01 ES024745-06 (to M.C.H), R01ES033462-0 (M.C.H). We acknowledge the shared facilities of the preclinical Imaging and Microscopy Resource and National Cancer Institute Cancer Center Support Grant 5P30 CA023108-37 at the Norris Cotton Cancer Center at Dartmouth, and NIH Grant P20-GM113132 to support the Institute for Biomolecular Targeting at Dartmouth. The Brain Luminex experiment was carried out in DartLab, the Immune Monitoring and Flow Cytometry Shared Resource at the Norris Cotton Cancer Center at Dartmouth, with NCI Cancer Center Support, Grant 5P30CA023108-41.

## Declarations

All the experiments were performed in accordance with legal and institutional guidelines and were carried out under ethics, consent and permissions of the Ethical Committee of Care and Use of Laboratory Animals at Geisel School of Medicine at Dartmouth.

## Consent for publication

All authors read and approved the manuscript.

## Availability of data and materials

The authors declares that the relevant data are included in the article.

## Competing Interests

The authors declared that they are no competing interests.

## Notes

### Competing Interest Statement

The authors have declared no competing interest.

## References

1. Reitz, C.; Rogaeva, E.; Beecham, G.W. Late-onset vs nonmendelian early-onset Alzheimer disease: A distinction without a difference? Neurol Genet 2020, 6, e512, doi:10.1212/nxg.0000000000000512.

2. Isik, A.T. Late onset Alzheimer’s disease in older people. Clin Interv Aging 2010, 5, 307–311, doi:10.2147/cia.S11718.

3. 2024 Alzheimer’s disease facts and figures. Alzheimer’s & Dementia 2024, 20, 3708–3821, doi:10.1002/alz.13809.

4. Fortea, J.; Pegueroles, J.; Alcolea, D.; Belbin, O.; Dols-Icardo, O.; Vaqué-Alcázar, L.; Videla, L.; Gispert, J.D.; Suárez-Calvet, M.; Johnson, S.C.; et al. APOE4 homozygozity represents a distinct genetic form of Alzheimer’s disease. Nat Med 2024, 30, 1284–1291, doi:10.1038/s41591-024-02931-w.

5. Liu, C.C.; Liu, C.C.; Kanekiyo, T.; Xu, H.; Bu, G. Apolipoprotein E and Alzheimer disease: risk, mechanisms and therapy. Nat Rev Neurol 2013, 9, 106–118, doi:10.1038/nrneurol.2012.263.

6. Steele, O.G.; Stuart, A.C.; Minkley, L.; Shaw, K.; Bonnar, O.; Anderle, S.; Penn, A.C.; Rusted, J.; Serpell, L.; Hall, C.; et al. A multi-hit hypothesis for an APOE4-dependent pathophysiological state. Eur J Neurosci 2022, 56, 5476–5515, doi:10.1111/ejn.15685.

7. Lin, Y.T.; Seo, J.; Gao, F.; Feldman, H.M.; Wen, H.L.; Penney, J.; Cam, H.P.; Gjoneska, E.; Raja, W.K.; Cheng, J.; et al. APOE4 Causes Widespread Molecular and Cellular Alterations Associated with Alzheimer’s Disease Phenotypes in Human iPSC-Derived Brain Cell Types. Neuron 2018, 98, 1141–1154 e1147, doi:10.1016/j.neuron.2018.05.008.

8. Tcw, J.; Qian, L.; Pipalia, N.H.; Chao, M.J.; Liang, S.A.; Shi, Y.; Jain, B.R.; Bertelsen, S.E.; Kapoor, M.; Marcora, E.; et al. Cholesterol and matrisome pathways dysregulated in astrocytes and microglia. Cell 2022, 185, 2213–2233 e2225, doi:10.1016/j.cell.2022.05.017.

9. Blanchard, J.W.; Akay, L.A.; Davila-Velderrain, J.; von Maydell, D.; Mathys, H.; Davidson, S.M.; Effenberger, A.; Chen, C.Y.; Maner-Smith, K.; Hajjar, I.; et al. APOE4 impairs myelination via cholesterol dysregulation in oligodendrocytes. Nature 2022, 611, 769–779, doi:10.1038/s41586-022-05439-w.

10. Machlovi, S.I.; Neuner, S.M.; Hemmer, B.M.; Khan, R.; Liu, Y.; Huang, M.; Zhu, J.D.; Castellano, J.M.; Cai, D.; Marcora, E.; et al. APOE4 confers transcriptomic and functional alterations to primary mouse microglia. Neurobiol Dis 2022, 164, 105615, doi:10.1016/j.nbd.2022.105615.

11. Litvinchuk, A.; Suh, J.H.; Guo, J.L.; Lin, K.; Davis, S.S.; Bien-Ly, N.; Tycksen, E.; Tabor, G.T.; Remolina Serrano, J.; Manis, M.; et al. Amelioration of Tau and ApoE4-linked glial lipid accumulation and neurodegeneration with an LXR agonist. Neuron 2024, 112, 384–403 e388, doi:10.1016/j.neuron.2023.10.023.

12. Haney, M.S.; Palovics, R.; Munson, C.N.; Long, C.; Johansson, P.K.; Yip, O.; Dong, W.; Rawat, E.; West, E.; Schlachetzki, J.C.M.; et al. APOE4/4 is linked to damaging lipid droplets in Alzheimer’s disease microglia. Nature 2024, 628, 154–161, doi:10.1038/s41586-024-07185-7.

13. Hippius, H.; Neundörfer, G. The discovery of Alzheimer’s disease. Dialogues Clin Neurosci 2003, 5, 101–108, doi:10.31887/DCNS.2003.5.1/hhippius.

14. Long, J.M.; Holtzman, D.M. Alzheimer Disease: An Update on Pathobiology and Treatment Strategies. Cell 2019, 179, 312–339, doi:10.1016/j.cell.2019.09.001.

15. Heneka, M.T.; Carson, M.J.; El Khoury, J.; Landreth, G.E.; Brosseron, F.; Feinstein, D.L.; Jacobs, A.H.; Wyss-Coray, T.; Vitorica, J.; Ransohoff, R.M.; et al. Neuroinflammation in Alzheimer’s disease. Lancet Neurol 2015, 14, 388–405, doi:10.1016/S1474-4422(15)70016-5.

16. Chan, R.B.; Oliveira, T.G.; Cortes, E.P.; Honig, L.S.; Duff, K.E.; Small, S.A.; Wenk, M.R.; Shui, G.; Di Paolo, G. Comparative lipidomic analysis of mouse and human brain with Alzheimer disease. J Biol Chem 2012, 287, 2678–2688, doi:10.1074/jbc.M111.274142.

17. Yang, L.G.; March, Z.M.; Stephenson, R.A.; Narayan, P.S. Apolipoprotein E in lipid metabolism and neurodegenerative disease. Trends in Endocrinology & Metabolism 2023, 34, 430–445, doi:10.1016/j.tem.2023.05.002.

18. Honig, L.S.; Barakos, J.; Dhadda, S.; Kanekiyo, M.; Reyderman, L.; Irizarry, M.; Kramer, L.D.; Swanson, C.J.; Sabbagh, M. ARIA in patients treated with lecanemab (BAN2401) in a phase 2 study in early Alzheimer’s disease. Alzheimers Dement (N Y) 2023, 9, e12377, doi:10.1002/trc2.12377.

19. Foley, K.E.; Wilcock, D.M. Three major effects of APOE(epsilon4) on Abeta immunotherapy induced ARIA. Front Aging Neurosci 2024, 16, 1412006, doi:10.3389/fnagi.2024.1412006.

20. Yen, J.-H.J.; Yu, I.-C.I. The role of ApoE-mediated microglial lipid metabolism in brain aging and disease. Immunometabolism 2023, 5, e00018, doi:10.1097/in9.0000000000000018.

21. Huttunen, H.J.; Kovacs, D.M. ACAT as a drug target for Alzheimer’s disease. Neurodegener Dis 2008, 5, 212–214, doi:10.1159/000113705.

22. Hutter-Paier, B.; Huttunen, H.J.; Puglielli, L.; Eckman, C.B.; Kim, D.Y.; Hofmeister, A.; Moir, R.D.; Domnitz, S.B.; Frosch, M.P.; Windisch, M.; et al. The ACAT inhibitor CP-113,818 markedly reduces amyloid pathology in a mouse model of Alzheimer’s disease. Neuron 2004, 44, 227–238, doi:10.1016/j.neuron.2004.08.043.

23. Puglielli, L.; Konopka, G.; Pack-Chung, E.; Ingano, L.A.; Berezovska, O.; Hyman, B.T.; Chang, T.Y.; Tanzi, R.E.; Kovacs, D.M. Acyl-coenzyme A: cholesterol acyltransferase modulates the generation of the amyloid beta-peptide. Nat Cell Biol 2001, 3, 905–912, doi:10.1038/ncb1001-905.

24. Murphy, S.R.; Chang, C.C.; Dogbevia, G.; Bryleva, E.Y.; Bowen, Z.; Hasan, M.T.; Chang, T.Y. Acat1 knockdown gene therapy decreases amyloid-β in a mouse model of Alzheimer’s disease. Mol Ther 2013, 21, 1497–1506, doi:10.1038/mt.2013.118.

25. Bryleva, E.Y.; Rogers, M.A.; Chang, C.C.; Buen, F.; Harris, B.T.; Rousselet, E.; Seidah, N.G.; Oddo, S.; LaFerla, F.M.; Spencer, T.A.; et al. ACAT1 gene ablation increases 24(S)-hydroxycholesterol content in the brain and ameliorates amyloid pathology in mice with AD. Proc Natl Acad Sci U S A 2010, 107, 3081–3086, doi:10.1073/pnas.0913828107.

26. Li, H.; Huynh, T.N.; Duong, M.T.; Gow, J.G.; Chang, C.C.Y.; Chang, T.Y. ACAT1/SOAT1 Blockade Suppresses LPS-Mediated Neuroinflammation by Modulating the Fate of Toll-like Receptor 4 in Microglia. Int J Mol Sci 2023, 24, doi:10.3390/ijms24065616.

27. Valencia-Olvera, A.C.; Balu, D.; Faulk, N.; Amiridis, A.; Wang, Y.; Pham, C.; Avila-Munoz, E.; York, J.M.; Thatcher, G.R.J.; LaDu, M.J. Inhibition of ACAT as a Therapeutic Target for Alzheimer’s Disease Is Independent of ApoE4 Lipidation. Neurotherapeutics 2023, 20, 1120–1137, doi:10.1007/s13311-023-01375-3.

28. De La Torre, A.L.; Huynh, T.N.; Chang, C.C.Y.; Pooler, D.B.; Ness, D.B.; Lewis, L.D.; Pannem, S.; Feng, Y.; Samkoe, K.S.; Hickey, W.F.; et al. Stealth Liposomes Encapsulating a Potent ACAT1/SOAT1 Inhibitor F12511: Pharmacokinetic, Biodistribution, and Toxicity Studies in Wild-Type Mice and Efficacy Studies in Triple Transgenic Alzheimer’s Disease Mice. Int J Mol Sci 2023, 24, doi:10.3390/ijms241311013.

29. Huynh, T.N.; Havrda, M.C.; Zanazzi, G.J.; Chang, C.C.Y.; Chang, T.Y. Inhibiting the Cholesterol Storage Enzyme ACAT1/SOAT1 in Myelin Debris-Treated Microglial Cell Lines Activates the Gene Expression of Cholesterol Efflux Transporter ABCA1. Biomolecules 2024, 14, 1301.

30. Melton, E.M.; Li, H.; Benson, J.; Sohn, P.; Huang, L.H.; Song, B.L.; Li, B.L.; Chang, C.C.Y.; Chang, T.Y. Myeloid Acat1/Soat1 KO attenuates pro-inflammatory responses in macrophages and protects against atherosclerosis in a model of advanced lesions. J Biol Chem 2019, 294, 15836–15849, doi:10.1074/jbc.RA119.010564.

31. Huang, L.H.; Melton, E.M.; Li, H.; Sohn, P.; Jung, D.; Tsai, C.Y.; Ma, T.; Sano, H.; Ha, H.; Friedline, R.H.; et al. Myeloid-specific Acat1 ablation attenuates inflammatory responses in macrophages, improves insulin sensitivity, and suppresses diet-induced obesity. Am J Physiol Endocrinol Metab 2018, 315, E340–E356, doi:10.1152/ajpendo.00174.2017.

32. Bohlen, C.J.; Bennett, F.C.; Tucker, A.F.; Collins, H.Y.; Mulinyawe, S.B.; Barres, B.A. Diverse Requirements for Microglial Survival, Specification, and Function Revealed by Defined-Medium Cultures. Neuron 2017, 94, 759-773.e758, doi:10.1016/j.neuron.2017.04.043.

33. Greenspan, P.; Fowler, S.D. Spectrofluorometric studies of the lipid probe, nile red. J Lipid Res 1985, 26, 781–789.

34. Cadigan, K.M.; Chang, C.C.; Chang, T.Y. Isolation of Chinese hamster ovary cell lines expressing human acyl-coenzyme A/cholesterol acyltransferase activity. J Cell Biol 1989, 108, 2201–2210, doi:10.1083/jcb.108.6.2201.

35. Chang, C.C.; Sakashita, N.; Ornvold, K.; Lee, O.; Chang, E.T.; Dong, R.; Lin, S.; Lee, C.Y.; Strom, S.C.; Kashyap, R.; et al. Immunological quantitation and localization of ACAT-1 and ACAT-2 in human liver and small intestine. J Biol Chem 2000, 275, 28083–28092, doi:10.1074/jbc.M003927200.

36. Rawat, V.; Wang, S.; Sima, J.; Bar, R.; Liraz, O.; Gundimeda, U.; Parekh, T.; Chan, J.; Johansson, J.O.; Tang, C.; et al. ApoE4 Alters ABCA1 Membrane Trafficking in Astrocytes. J Neurosci 2019, 39, 9611–9622, doi:10.1523/jneurosci.1400-19.2019.

37. Varshney, P.; Yadav, V.; Saini, N. Lipid rafts in immune signalling: current progress and future perspective. Immunology 2016, 149, 13–24, doi:10.1111/imm.12617.

38. Zhou, Y.; Chen, Y.; Xu, C.; Zhang, H.; Lin, C. TLR4 Targeting as a Promising Therapeutic Strategy for Alzheimer Disease Treatment. Front Neurosci 2020, 14, 602508, doi:10.3389/fnins.2020.602508.

39. Davies, D.A. The Role of APOE and NF-κB in Alzheimer’s Disease. Immuno 2021, 1, 391–399.

40. Ophir, G.; Amariglio, N.; Jacob-Hirsch, J.; Elkon, R.; Rechavi, G.; Michaelson, D.M. Apolipoprotein E4 enhances brain inflammation by modulation of the NF-kappaB signaling cascade. Neurobiol Dis 2005, 20, 709–718, doi:10.1016/j.nbd.2005.05.002.

41. Tai, L.M.; Ghura, S.; Koster, K.P.; Liakaite, V.; Maienschein-Cline, M.; Kanabar, P.; Collins, N.; Ben-Aissa, M.; Lei, A.Z.; Bahroos, N. APOE-modulated Aβ-induced neuroinflammation in Alzheimer’s disease: current landscape, novel data, and future perspective. Journal of neurochemistry 2015, 133, 465–488.

42. Meier-Soelch, J.; Mayr-Buro, C.; Juli, J.; Leib, L.; Linne, U.; Dreute, J.; Papantonis, A.; Schmitz, M.L.; Kracht, M. Monitoring the Levels of Cellular NF-κB Activation States. Cancers (Basel) 2021, 13, doi:10.3390/cancers13215351.

43. De La Torre, A.L.; Smith, C.; Granger, J.; Anderson, F.L.; Harned, T.C.; Havrda, M.C.; Chang, C.C.Y.; Chang, T.Y. Facile method to incorporate high-affinity ACAT/SOAT1 inhibitor F12511 into stealth liposome-based nanoparticle and demonstration of its efficacy in blocking cholesteryl ester biosynthesis without overt toxicity in neuronal cell culture. J Neurosci Methods 2022, 367, 109437, doi:10.1016/j.jneumeth.2021.109437.

44. Mhatre-Winters, I.; Eid, A.; Han, Y.; Tieu, K.; Richardson, J.R. Sex and APOE Genotype Alter the Basal and Induced Inflammatory States of Primary Microglia from APOE Targeted Replacement Mice. Int J Mol Sci 2022, 23, doi:10.3390/ijms23179829.

45. Mishra, A.; Soto, M.; Delatorre, N.; Rodgers, K.E.; Brinton, R.D. APOE4 genetic burden and female sex impact immune profile in brain and periphery in aged mice. Alzheimer’s & Dementia 2021, 17, e056541, doi:10.1002/alz.056541.

46. Kawahara, K.; Suenobu, M.; Yoshida, A.; Koga, K.; Hyodo, A.; Ohtsuka, H.; Kuniyasu, A.; Tamamaki, N.; Sugimoto, Y.; Nakayama, H. Intracerebral microinjection of interleukin-4/interleukin-13 reduces β-amyloid accumulation in the ipsilateral side and improves cognitive deficits in young amyloid precursor protein 23 mice. Neuroscience 2012, 207, 243–260, doi:10.1016/j.neuroscience.2012.01.049.

47. Beklen, A.; Sarp, A.S.; Uckan, D.; Tsaous Memet, G. The function of TLR4 in interferon gamma or interleukin-13 exposed and lipopolysaccharide stimulated gingival epithelial cell cultures. Biotechnic & Histochemistry 2014, 89, 505–512, doi:10.3109/10520295.2014.903299.

48. Montagne, A.; Nation, D.A.; Sagare, A.P.; Barisano, G.; Sweeney, M.D.; Chakhoyan, A.; Pachicano, M.; Joe, E.; Nelson, A.R.; D’Orazio, L.M.; et al. APOE4 leads to blood-brain barrier dysfunction predicting cognitive decline. Nature 2020, 581, 71–76, doi:10.1038/s41586-020-2247-3.

49. Gouna, G.; Klose, C.; Bosch-Queralt, M.; Liu, L.; Gokce, O.; Schifferer, M.; Cantuti-Castelvetri, L.; Simons, M. TREM2-dependent lipid droplet biogenesis in phagocytes is required for remyelination. J Exp Med 2021, 218, doi:10.1084/jem.20210227.

50. Rogers, M.A.; Chang, C.C.Y.; Maue, R.A.; Melton, E.M.; Peden, A.A.; Garver, W.S.; Lee, J.; Schroen, P.; Huang, M.; Chang, T.Y. Acat1/Soat1 knockout extends the mutant Npc1 mouse lifespan and ameliorates functional deficiencies in multiple organelles of mutant cells. Proc Natl Acad Sci U S A 2022, 119, e2201646119, doi:10.1073/pnas.2201646119.

51. Lee, S.; Devanney, N.A.; Golden, L.R.; Smith, C.T.; Schwartz, J.L.; Walsh, A.E.; Clarke, H.A.; Goulding, D.S.; Allenger, E.J.; Morillo-Segovia, G.; et al. APOE modulates microglial immunometabolism in response to age, amyloid pathology, and inflammatory challenge. Cell Rep 2023, 42, 112196, doi:10.1016/j.celrep.2023.112196.

52. Tan, W.; Zhang, Q.; Dong, Z.; Yan, Y.; Fu, Y.; Liu, X.; Zhao, B.; Duan, X. Phosphatidylcholine Ameliorates LPS-Induced Systemic Inflammation and Cognitive Impairments via Mediating the Gut-Brain Axis Balance. J Agric Food Chem 2020, 68, 14884–14895, doi:10.1021/acs.jafc.0c06383.

53. Pathak, P.; London, E. Unsaturated Phosphatidylcholine Acyl Chain Structure Affects the Size of Ordered Nanodomains (Lipid Rafts) Formed by Sphingomyelin and Cholesterol. Biophysical Journal 2009, 96, 363a, doi:10.1016/j.bpj.2008.12.1955.

54. Xian, X.; Pohlkamp, T.; Durakoglugil, M.S.; Wong, C.H.; Beck, J.K.; Lane-Donovan, C.; Plattner, F.; Herz, J. Reversal of ApoE4-induced recycling block as a novel prevention approach for Alzheimer’s disease. eLife 2018, 7, doi:10.7554/eLife.40048.

55. Victor, M.B.; Leary, N.; Luna, X.; Meharena, H.S.; Scannail, A.N.; Bozzelli, P.L.; Samaan, G.; Murdock, M.H.; von Maydell, D.; Effenberger, A.H.; et al. Lipid accumulation induced by APOE4 impairs microglial surveillance of neuronal-network activity. Cell Stem Cell 2022, 29, 1197-1212.e1198, doi:10.1016/j.stem.2022.07.005.

56. Scheiblich, H.; Dansokho, C.; Mercan, D.; Schmidt, S.V.; Bousset, L.; Wischhof, L.; Eikens, F.; Odainic, A.; Spitzer, J.; Griep, A.; et al. Microglia jointly degrade fibrillar alpha-synuclein cargo by distribution through tunneling nanotubes. Cell 2021, 184, 5089-5106.e5021, doi:10.1016/j.cell.2021.09.007.

57. Adomshick, V.; Pu, Y.; Veiga-Lopez, A. Automated lipid droplet quantification system for phenotypic analysis of adipocytes using CellProfiler. Toxicol Mech Methods 2020, 30, 378–387, doi:10.1080/15376516.2020.1747124.

